# Mitochondrial genome editing reveals the impact of complex I heteroplasmic variants on human neurogenesis

**DOI:** 10.64898/2026.10.28.762518

**Authors:** Isabella Tolle, Claudio Fiorini, Jochen Dobner, Annika Zink, Alberto Pietro Pasti, Valeria Balmaceda, Florian Tschuschke, Diran Herebian, Ertan Mayatepek, Emanuela Bottani, Pawel Lisowski, Valerio Carelli, Andrea Rossi, Alessandro Prigione

**Affiliations:** Department of General Pediatrics, Neonatology and Pediatric Cardiology, Medical Faculty and University Hospital Düsseldorf, Heinrich Heine University Düsseldorf, Düsseldorf, Germany; IRCCS Istituto delle Scienze Neurologiche di Bologna, Programma Neurogenetica, Bologna, Italy; IUF-Leibniz Research Institute for Environmental Medicine, Düsseldorf, Germany; Department of Diagnostics and Public Health, University of Verona, 37134 Verona, Italy; Berlin Institute for Medical Systems Biology (BIMSB), Max Delbrück Center for Molecular Medicine in the Helmholtz Association (MDC), 10115 Berlin, Germany; Department of Molecular Biology, Institute of Genetics and Animal Biotechnology, Polish Academy of Sciences, 05-552 Jastrzebiec n/Warsaw, Poland; Department of Biomedical and Neuromotor Sciences, University of Bologna, Bologna, Italy

**Author notes:** CORRESPONDENCE: Alessandro Prigione, MD, PhD. Heinrich Heine University (HHU), Medical Faculty, Department of General Pediatrics, Neonatology and Pediatric Cardiology, Moorenstr. 5, 40225 Düsseldorf, Germany.

**Keywords:** Mitochondrial diseases, Leigh syndrome, mtDNA, base editing, mitochondrial genome engineering, DdCBE, heteroplasmy, iPSCs

## Abstract

Mitochondrial DNA (mtDNA) variants, whether maternally inherited or acquired somatically during aging, can be particularly detrimental to cells that rely on mitochondrial metabolism such as neurons. The functional consequences of mtDNA variants are thought to depend on their heteroplasmy, defined as the proportion of mtDNA molecules carrying that variant. However, investigating the specific effects of heteroplasmic mtDNA variants on human neuronal function has been challenging. Studies using patient-derived cells cannot conclusively attribute observed phenotypes to a single mtDNA variant, while conventional genome editing approaches are not applicable to mtDNA. Here, we use DddA-derived cytosine base editor (DdCBE) to introduce pathogenic heteroplasmic mtDNA variants into Neurogenin 2 (NGN2)-inducible human induced pluripotent stem cells (iPSCs). We focus on two variants in the genes *MT-ND1* and *MT-ND5* belonging to complex I (CI), the largest component of the mitochondrial respiratory chain and a frequent contributor to human disease, including the incurable neurodevelopmental disorder Leigh syndrome. mtDNA-edited iPSC lines exhibit low off-target editing and maintain similar heteroplasmy following subclonal isolation and differentiation into NGN2-induced neurons (iNs). Although the CI variants do not substantially affect mitochondrial energy metabolism in undifferentiated iPSCs, they reduce mitochondrial membrane potential in iNs where they drive a transcriptional reprogramming indicative of impaired neuronal commitment. These findings suggest that heteroplasmic CI variants are sufficient to disrupt neural fate specification even at relatively low heteroplasmy. More broadly, mtDNA editing in human iPSCs provides a powerful platform to investigate how mitochondrial fitness and heteroplasmy influence development and disease.

## INTRODUCTION

Mitochondria are double-membrane organelles carrying their own genome that execute key cellular functions, including bioenergetics, redox and calcium homeostasis, and signal transduction.^1^ Mitochondrial DNA (mtDNA) is a small circular double-stranded molecule of approximately 16.6 kb size encoding only 37 genes, among which 13 code for proteins.^2^ All of these essential proteins are distributed throughout 4 of the 5 respiratory chain complexes, with 7 proteins belonging to complex I (CI).^3^ In the mitochondrial respiratory chain, CI is the first and largest component and is viewed as the rate-limiting step in oxidative phosphorylation (OXPHOS).^4,5^ CI defects are the first cause of primary mitochondrial diseases, monogenic disorders that disrupt OXPHOS and particularly affect tissues with high energy requirements such as the central nervous system (CNS) and the cardiac and skeletal muscle.^6–9^ A severe form of mitochondrial disease is Leigh syndrome (LS, OMIM #256000), which causes progressive retardation of psychomotor function and early death.^10–12^ Pathogenic variants in the mtDNA genes encoding for CI proteins could lead to LS,^13^ including m.3890G>A in the gene *MT-ND1*^14^ or m.13513G>A in the gene *MT-ND5*^*15*^ (Figure 1A).

**Figure 1.**
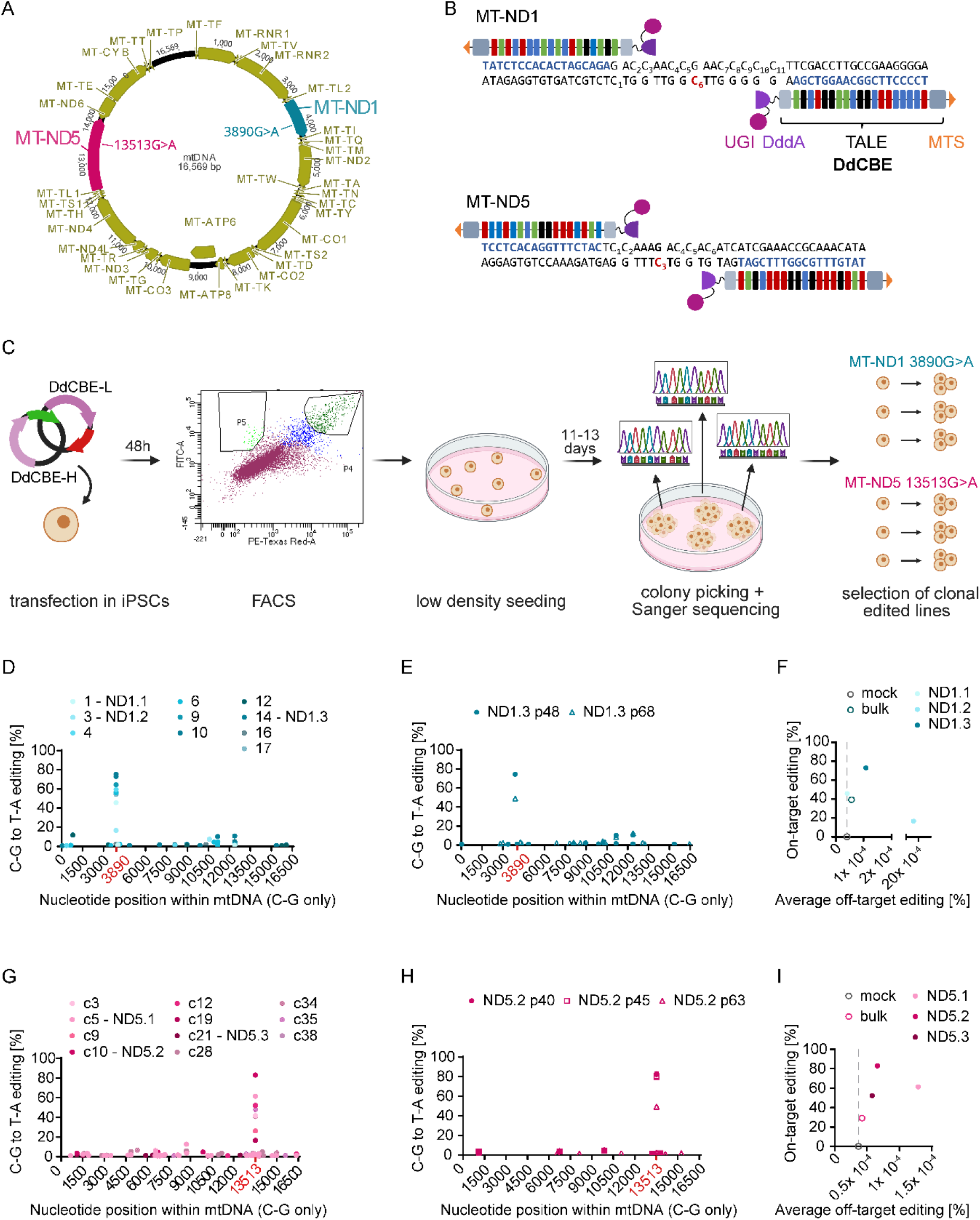
Introducing CI heteroplasmic variants in human iPSCs via mtDNA base editing. (A) Schematic of human mtDNA with highlighted targeted variants in CI genes MT-ND1 (teal) and MT-ND5 (magenta). (B) Schematic of DdCBE architecture at MT-ND1 and MT-ND5 target sites. TALE-binding sites are highlighted in blue, targeted cytosines are highlighted in red, underscoring numbers indicate cytosines in the spacing region. UGI: uracil glycosylase inhibitor, DddA: double-stranded DNA deaminase toxin A, TALE: transcription activator-like effector, MTS: mitochondrial targeting signal. (C) Schematic of mtDNA editing workflow in iPSCs. The most efficient DdCBE pairs were transfected into iPSCs and double-positive transfectants were sorted via FACS. After low density seeding, single colonies were picked and screened via Sanger sequencing. For each target site, three successfully edited colonies were chosen for downstream experiments. (D) NGS of whole mtDNA of all MT-ND1 edited iPSC colonies that were identified by Sanger sequencing. (E) Editing stability of ND1.3 iPSC line upon long-term cultivation. (F) On-target editing vs average off-target editing of mock-treated control cells, bulk edited cells, and three selected edited lines (ND1.1, ND1.2, ND1.3). (G) NGS of whole mtDNA of all MT-ND5 edited iPSC colonies that were identified by Sanger sequencing. (H) Editing stability of ND5.2 iPSC line upon long-term cultivation. (I) On-target editing vs average off-target editing of mock-treated control cells, bulk edited cells, and three selected edited lines (ND5.1, ND5.2, ND5.3).

As mammalian cells contain multiple copies of mtDNA, pathogenic variants can affect a variable proportion of mtDNA molecules. This proportion, defined as heteroplasmy, is believed to dictate their functional impact.^16,17^ mtDNA variants are generally considered functionally recessive, as they must be present within a significant amount of mtDNA molecules to cause biochemical impairment. Higher heteroplasmy is usually associated with more severe conditions. However, this association is not entirely linear, as similar heteroplasmic variants can cause different disease entities.^18^ m.13513G>A has been even suggested to act dominantly, as it can cause functional defects at low heterosplasmy.^19^ The potential pathogenicity at relatively low levels may be specific to CI variants.^11,12,19^ In fact, complex V (CV) mtDNA variants are typically present in almost all mtDNA copies in LS individuals, as lower levels of heteroplasmic CV variants are instead associated with a milder disease presentation called neuropathy, ataxia, and retinitis pigmentosa (NARP).^20^

The correlation of heteroplasmic mtDNA variants to defined biochemical and cellular defects was established through transmitochondrial cytoplasmic hybrids (cybrids),^21,22^ produced by fusing enucleated donor cells obtained from patients with recipient immortalized cells devoid of mtDNA. These studies showed that increasing heteroplasmy leads to progressive changes in transcriptional response.^23^ However, cybrids are based on immortalized cancer-like cells, which mainly rely on glycolysis for energy production and do not exhibit the functional properties of neurons.^24^ The consequences of mtDNA variants in human neuronal cells have been addressed using differentiated induced pluripotent stem cells (iPSCs) obtained from affected patients.^18,25,26^ As the level of heteroplasmy can vary among iPSC lines derived from the same somatic cells, reprogramming patient-derived cells enables the generation of iPSC lines carrying different degrees of heteroplasmy from the same patient cells.^27–29^ Nonetheless, studies using patient-derived cells cannot conclude that the observed phenotypes stem solely from the presence of a given mtDNA variant. In fact, it is known that individuals carrying the same mtDNA variants might show distinct symptomatology possibly due to nuclear background contribution, as specific nuclear genetic loci might act as modifiers of pathogenic mtDNA variants.^17,30,31^

Here, we aim to demonstrate whether the sole presence of heteroplasmic CI variants can lead to functional defects and transcriptional reprogramming in healthy neuronal cells. Precise manipulation of mtDNA has been historically difficult due to the polyploid nature of mtDNA, inefficient nucleic acid transport into mitochondria, as well as the lack of efficient DNA recombination.^32^ These features hinder the engineering of mtDNA by classical methods like clustered regularly interspaced short palindromic repeats/CRISPR-associated protein (CRISPR/Cas).^33^ Hence, we took advantage of novel mtDNA base editing approaches based on CRISPR-free double-stranded DNA deaminase toxin A (DddA)-derived cytosine base editor (DdCBE).^34^ While CRISPR/Cas-based editing approaches rely on guide RNA to target the genomic region of interest, DdCBE makes use of transcription activator-like effectors (TALEs), proteins that can be programmed to specifically bind a chosen DNA sequence and can be targeted for mitochondrial import through fusion with a mitochondrial targeting sequence (MTS). The distinguishing feature of DdCBE compared to previous base editors, is the DNA-modifying component, the cytidine deaminase DddA, which in contrast to its predecessors is capable of deaminating double-stranded DNA and thus does not rely on proteins such as Cas9 to melt the double strand for access to the target base.^34,35^

We applied DdCBE to introduce two heteroplasmic CI variants associated with LS (m.3890G>A in the gene *MT-ND1* and m.13513G>A in the gene *MT-ND5*) into wildtype (WT) human iPSCs that we previously engineered to carry an inducible cassette for Neurogenin 2 (NGN2), which enables the rapid generation of (NGN2), which enables the rapid generation of induced neurons (iNs).^36^ We found that CI heteroplasmic variants could be effectively introduced into WT iPSCs without substantial off-target editing. Heteroplasmy levels decreased during long-term expansion of iPSCs but remained similar upon subclonal passaging and conversion into iNs. The consequences of heteroplasmy differed between iPSCs and iNs. While CI variants did not affect the metabolic properties of iPSCs, in iNs they caused a decrease of mitochondrial membrane potential (MMP) paralleled by a transcriptional reconfiguration suggestive of impaired neuronal fate commitment even at relatively low levels of heteroplasmy.

The results imply that germline inherited CI mtDNA variants could lead to functional defects during tissue specification. DdCBE has previously been used to manipulate mtDNA in immortalized and primary human somatic cells,^34^ in human embryos,^37,38^ to generate mtDNA disease models in mice,^39,40^ rats,^41,42^ and zebrafish,^43^ as well as in a proof of concept study to correct a pathological mutation in iPSCs.^44^ To our knowledge, however, this is the first report of the targeted installation of a pathogenic mtDNA variant into a healthy human iPSC line to generate an isogenic disease model. Such an approach opens the way to precise disease modeling of primary mitochondrial diseases and may shed light on the currently unknown mechanisms underlying the maintenance and tissue-specific distribution and segregation of mtDNA heteroplasmy during development and disease.

## RESULTS

### Precise introduction of LS-associated CI mtDNA variants in human iPSCs

We first identified two LS-associated CI mtDNA variants amenable to mtDNA base editing. As wildtype DddA can only enable conversion of C•G base pair (bp) to T•A bp through deamination of cytidines at T**C** (or T**CC**) sites in the mtDNA,^34^ we selected two CI variants that could be targeted: m.3890G>A (p.Arg195Gln) in the gene *MT-ND1*^14,45^ and m.13513G>A (p.Asp393Asn) in the gene *MT-ND5*^*15*^ (Figure 1A-B). DddA is split into two halves to avoid widespread deamination, whereby two different split variants exist. For guiding the two halves to the target locus where DddA is reconstituted and can deaminate double-stranded DNA, DdCBE does not depend on gRNAs, which cannot cross the mitochondrial double membrane, but on TALEs proteins designed to bind upstream (on the light strand, L) and downstream (on the heavy strand, H) of the target site.^34^ The DdCBE constructs are targeted to mitochondria by MTS and a copy of uracil glycosylase inhibitor (UGI) on each half prevents the removal of the newly installed mutation through base excision repair (Figure 1B). Because the choice of DddA split variant (G1333 or G1397) and the arrangement of the DddA halves relative to the L and H strands dictate the editing success depending on the architecture of the target site,^34^ we screened all possible orientations in HEK cells to determine the most effective DdCBE pairs (Figure S1A). We then applied a pair with high on-target and no bystander editing (L-1333-C + H-1333-N for both targets) to WT iPSCs. We included fluorescence markers mCherry and Neongreen to the two DdCBE halves to allow sorting cells that had taken up both constructs (Figure 1C). After low density seeding and single colonies picking, we used Sanger sequencing to identify successfully edited clones (Figure 1C). The edited clones were then subjected to next generation sequencing (NGS) of whole mtDNA (Figure 1D, Figure 1G) and compared to mock-treated controls, where only the H-strand DdCBE half was transfected. Calls that were identified in the mock-treated controls were deemed to be editing-independent SNVs and removed from the graphs. NGS revealed that editing efficiencies ranged from 3 to 75 % for the *MT-ND1* target site, with most lines centering around roughly 50 %, and 17-83 % for the *MT-ND5* site with a more diverse distribution of editing efficiencies (Figure 1D, Figure 1G). Throughout whole mtDNA only few off-target events were detected (1-12 for *MT-ND1* and 0-41 for *MT-ND5*) and a general trend of increasing number of off-target events with increasing on-target editing could be observed (Figure S1F). Most of off-target incidents occurred with a frequency of >3 % (Figure S1D, Figure S1E). Except for two instances at the *MT-ND5* site (>2 %), no bystander editing was detected (Table S1, Table S2). On-target editing was relatively stable over a few passages, but decreased upon long-term passaging (Figure 1E, Figure 1H). Similar results were obtained using our recently developed Nanopore-based mtDNA sequencing approach Mitopore^46^ (Figure S1B-C). In summary, we were able to obtain edited iPSC lines with high on-target editing, no bystander and low mtDNA-wide off-target editing for both target sites.

We selected three clones for each CI variant for downstream experiments: ND1.1, ND1.2, ND1.3 carrying the variant m.3890G>A in the gene *MT-ND1* and ND5.1, ND5.2, ND5.3 carrying the variant m.13513G>A in the gene *MT-ND5*. The selected iPSC lines carried the introduced CI variants at different levels of heteroplasmy and exhibited low amount of off-target editing (Figure 1F, Figure 1I). As the nuclear genome contains stray mtDNA fragments called nuclear mitochondrial DNA fragments (NUMTs), which may have a strong similarity to the mitochondrial target sites, we next screened for putative TALE binding sites in the nuclear genome.^47,48^ For each target site we chose the three regions with the highest binding affinity for both TALEs.^49^ No nuclear off-target editing could be detected in these regions in the six selected iPSC lines (Figure S2A-B). The six mtDNA edited iPSC lines also retained pluripotency features (Figure S3A-B) and a normal karyotype (Figure S4 and Figure S5).

A defined heteroplasmy value in cell lines composed of different individual cells may correspond to the heteroplasmy of each individual cell or may represent an average of different heteroplasmy values in the individual cells. To address this aspect, we performed subcloning of ND5.2 and ND5.3 lines (Figure 2A), which had heteroplasmy levels around 80 % and 50 % respectively (Figure 1I). NGS-based whole mtDNA sequencing showed that subclones generated from both lines retained comparable heteroplasmy (75-87 % and 41-58 %) to the respective original iPSC lines (Figure 2B-C). These results suggest that individual cells within the iPSC lines may carry the variant at similar heteroplasmy levels. Therefore, changes in heteroplasmy in the cell lines might reflect concomitant changes in the heteroplasmy of the individual cells rather than changes in the composition of the population.

**Figure 2.**
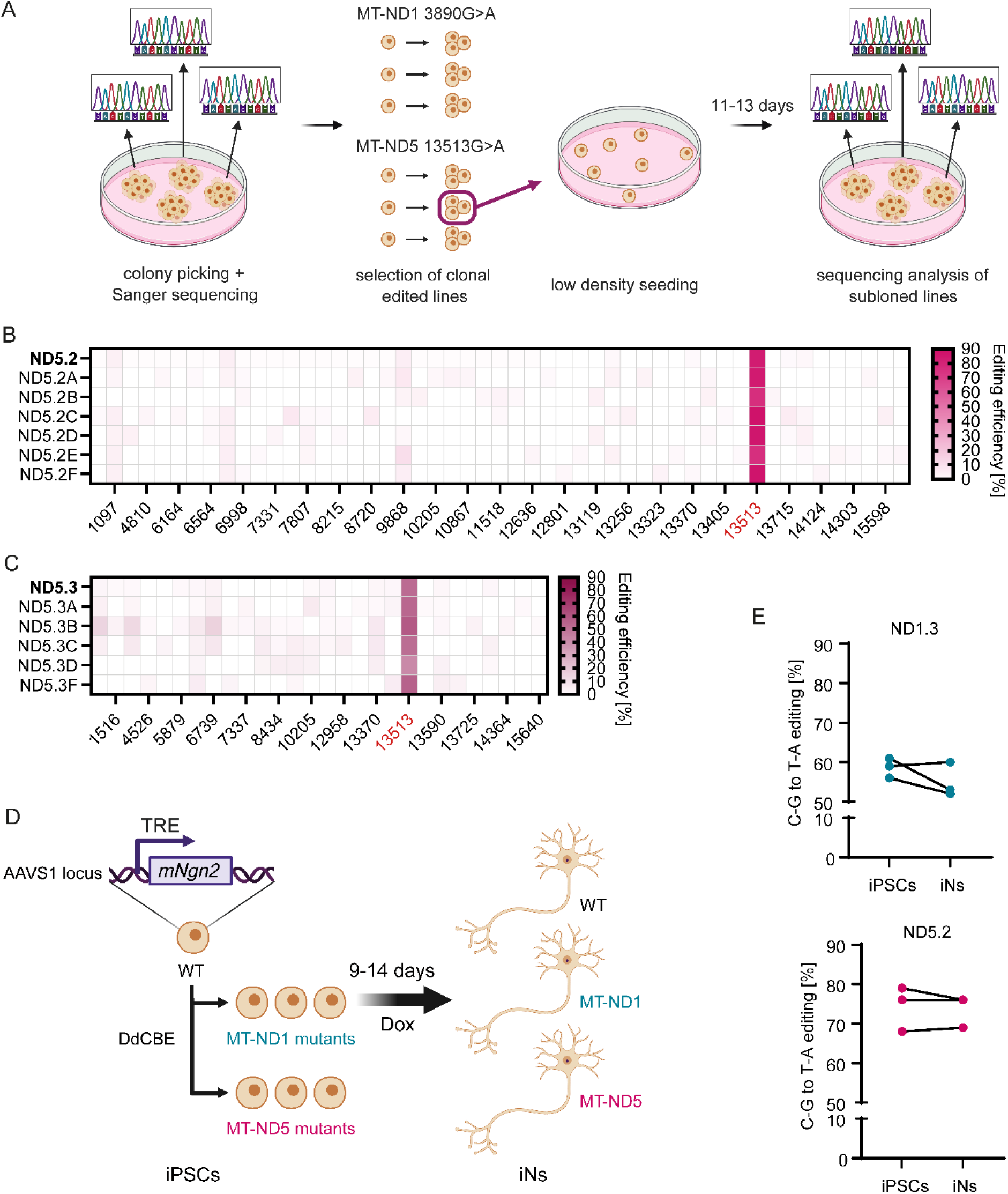
Heteroplasmy is maintained during subcloning and neuronal induction. (A) Schematic of subcloning workflow. Edited iPSC lines ND5.2, ND5.3 were singularized and seeded in low density and six or five single subclones were picked for each line. (B-C) NGS of whole mtDNA of the iPSC subclones obtained from ND5.2 and ND5.3. (D) The human iPSC line chosen for mtDNA base editing harbors an inducible Neurogenin2 (NGN2) expression cassette that allowed the rapid generation of induced neurons (iNs) upon doxycycline (Dox) exposure. (E) Heteroplasmy levels of the mtDNA edited lines ND1.3 (top) and ND5.2 (bottom) before and after differentiation of iPSCs into iNs.

### Heteroplasmic CI mtDNA variants disrupt neuronal fate commitment

We differentiated the mtDNA edited iPSC lines into induced neurons (iNs).^36^ Since the WT iPSC line used for editing carried an inducible NGN2 cassette, iNs could be rapidly obtained by exposing iPSCs to doxycycline, which triggers NGN2 expression thereby inducing the conversion into neurons (Figure 2D, Figure S7). iNs maintained similar heteroplasmy levels as their respective parental iPSC lines prior to differentiation (Figure 2E).

Next, we addressed the functional consequences of the introduced CI variants. Specific CI activity was measured using iPSC Lysates, revealing that both heteroplasmic CI variants caused a slight, but not significant reduction in CI activity (Figure 3A). This is in line with previous experiments in cybrids, where m.3890G>A in *MT-ND1* caused significant reduction in CI activity only when homoplasmic and resulted in a milder decrease when heteroplasmic.^45^ In contrast, for m.13513G>A in *MT-ND5* there seems to be no correlation between heteroplasmy and enzymatic CI activity in patients,^50,51^ possibly due to the spatial separation of the redox reactions and the proton pumping machinery of the complex.^52^ Whereas this assay measures the oxidation of NADH in the peripheral arm of CI, ND5 is a core subunit of the membrane arm involved in proton pumping.^53,54^ Therefore, despite exhibiting only a mild decrease in NADH/CoQ redox chemistry, the mutation might cause more prominent dysfunction further downstream of the electron transport chain. We then measured mitochondrial membrane potential (MMP) in undifferentiated iPSCs. MMP appeared to be not consistently disrupted in all mtDNA edited iPSCs (Figure 3B). ND5.2 exhibited no disruption, while ND1.3 showed significantly decreased MMP only in 2 out of 5 trials. Consistently, ATP amount or metabolites of TCA cycle and glycolysis were not altered in undifferentiated iPSCs (Figure S6A-C).

When we measured MMP in iNs, we found that all mtDNA edited iNs displayed a significant decrease of MMP (Figure 3I), consistent with a proton pumping defect. These findings imply that the impact of heteroplasmic CI variants is different depending on the cell type and that neuronal cells may be more susceptible to CI deficiency possibly due to their higher OXPHOS reliance.

To determine the transcriptional response of neurons carrying CI variants, we performed bulk RNA sequencing of mtDNA edited iNs and the respective isogenic WT iNs (Figure 3C-G). For each mtDNA edited line, we screened three individual differentiation experiments. Each resulting batch of iNs thus carried slightly different heteroplasmy depending on the level of the parental iPSCs. We deemed bulk transcriptomics to sufficiently reflect the transcriptional state of the cells, as the cultures were fairly homogenous both in terms of cell type as well as heteroplasmy; NGN2 overexpression forced the cells to convert to iNs and those that failed to commit were removed by cytosine arabinoside (AraC) treatment,^55,56^ moreover, heteroplasmy was retained at similar levels upon subcloning (Figure 2A-C) and differentiation (Figure 2E).

Across all iNs carrying CI variants, downregulated genes encoded for proteins involved in neuronal development (e.g. *NEUROD2, ISL2, SEMA3D, PHOX2B, ARC, CHGA, BRINP2*) synaptic function and neuronal firing (e.g. *SYT3, NKX6-1, SHISA9, KCNN1, STAC, COMT, CHAT, MAOA, TAC1, GRIN1, SYN3, PHF24*), transmembrane proteins expressed in the CNS (e.g. *TMEM255B, TMEM151A*), proteins involved in RNA regulation for CNS formation and mitochondrial integrity (e.g. *SMG8, PCBP1*), and oxygen and nitric oxide regulators (e.g. *CYGB*) (Figure 3C-G, Table S1). Upregulated genes comprised genes involved in cell adhesion (e.g. *CTNNA3, LAMA4, PCDHA5, RELN, ITGA8*) growth and proliferation (e.g. *BMP3, CCDC152, PDGFD, HAPLN1, ANKRD18A, MEF2C, ETV1, PLCD4*) metabolism and ion regulation (e.g. *GCNT4, PMEL, ATP1A2*) (Figure 3C-G, Table S1). A common signature of differentially expressed genes (DEG) across all iNs indicated that key genes were similarly modulated by the different CI heteroplasmic variants (Figure 3H, Table S1). We could not identify a clear association between the gene expression level of these common transcripts and the heteroplasmy of iNs, as the presence of relatively low levels of heteroplasmy was sufficient to elicit similar transcriptional reprogramming in neurons.

**Figure 4.**
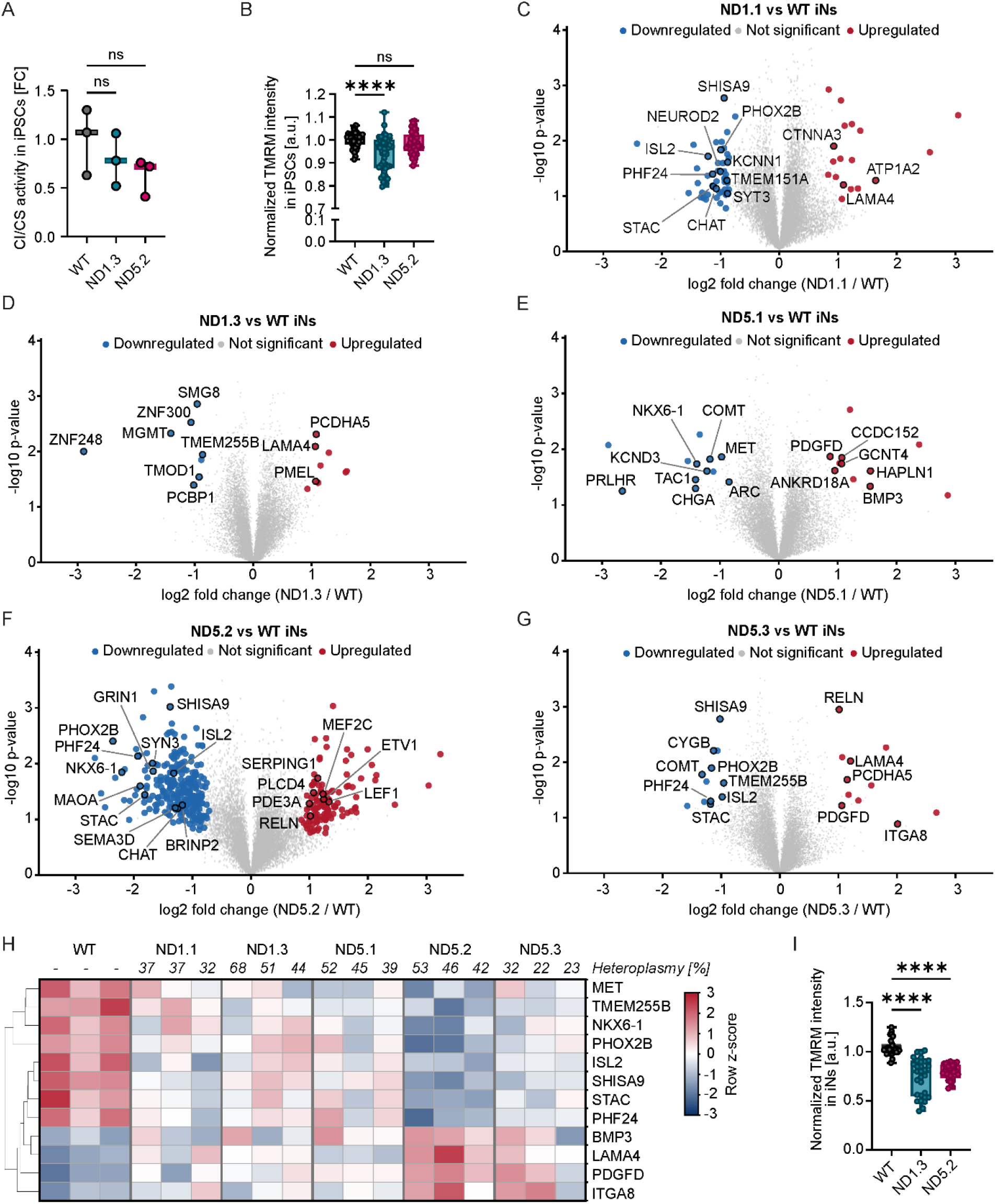
Heteroplasmic CI variants impair mitochondrial function and the neuronal commitment transcriptional program in iNs. (A) Complex I (CI) activity normalized over citrate synthase (CS) activity in mtDNA edited iPSC lines ND1.3 and ND5.2 compared to isogenic wildtype iPSCs (WT). Bars and boxes reflect the mean ± s.d. of n=3 independent biological replicates; ns: not significant; Brown-Forsythe and Welch ANOVA with Dunnett’s T3 multiple comparison. (B) High-content analysis (HCA)-based quantification of mitochondrial membrane potential (MMP) measured as TMRM intensity against Hoechst signal in iPSCs. Dots represent mean values per well with 14 images taken per well and 10 wells measured for each of n=5 independent experiments. **p<0.0001; Brown-Forsythe and Welch ANOVA with Dunnett’s T3 multiple comparison. (C-G) Volcano plots of differentially expressed genes (DEGs) in iNs derived from mtDNA edited iPSC lines ND1.1, ND1.3, ND5.1, ND5.2, and ND5.3 compared to isogenic wildtype iNs (WT). Significantly down- or up-regulated genes are highlighted in blue and red respectively. (H) Heat map of key significant DEGs shared between the five heteroplasmic mtDNA edited iNs compared to WT iNs. (I) HCA-based quantification of MMP measured as TMRM intensity against Hoechst signal in iNs. Dots represent mean values per well with 14 images taken per well and 10 wells measured for each of n=3 independent experiments. ****p<0.0001; Kruskal-Wallis test with Dunn’s multiple comparison.

## DISCUSSION

CI deficiency is the most frequent cause of mitochondrial dysfunction in humans, as it has been associated with common conditions such as diabetes and Parkinson’s disease.^6–8,57^ Indeed, CI defects are the first cause of mitochondrial diseases.^58,59^ Variants in the 7 CI mtDNA genes can cause LS, one of the most severe forms of mitochondrial disease.^13^ Heteroplasmic mtDNA variants are also observed in multiple tissues during aging and neurodegeneration.^16,60^ However, investigating the specific impact of mtDNA variants has been historically challenging.^61^ mtDNA cannot be targeted using CRIPSR/Cas-based genome editing.^33^ Furthermore, the use of patient-derived cellular models may not allow pinpointing the direct contribution of heteroplasmic mtDNA variants, as individual nuclear background may influence disease phenotype. Here, by employing DdCBE mtDNA base editing to introduce two CI variants in healthy human iPSCs, we were able to shed light on the specific functional impact of heteroplasmic CI variants in proliferating stem cells and derived neurons.

Similar heteroplasmy levels caused different functional impacts in different cell types. mtDNA edited neurons exhibited mitochondrial dysfunctions that were not observed in undifferentiated iPSCs. This could be because undifferentiated iPSCs mainly rely on glycolysis and then undergo a metabolic shift towards OXPHOS during neural fate commitment.^62–64^ Cell type identity and their specific metabolism could thus matter in explaining the phenotypes of mtDNA diseases. In fact, it is known that tissues relying on OXPHOS, such as the CNS, are particularly susceptible to mitochondrial dysfunction and that neurodegenerative diseases characteristically develop an energy gap.^9,65^ The lack of functional impact in undifferentiated iPSCs is in agreement with previous studies based on patient-derived iPSCs carrying mtDNA variants.^26,28,29,66–70^ mtDNA variants might therefore be retained within embryonic and adult stem cells because they are functionally irrelevant in these cells and would then cause dysfunctions within differentiated cells during tissue specification.

The transcriptional signature of neurons carrying CI mtDNA variants highlighted the disruption of the neurodevelopmental program. Dysregulated genes were involved in neurogenesis and neuronal maturation. These findings imply that CI-related mitochondrial diseases may display a neurodevelopmental component. In fact, it is known that mitochondrial fitness is crucial during development, where it influences the specific temporality of CNS formation.^71,72^ Accordingly, antenatal manifestations can be observed in LS and other mitochondrial diseases,^73,74^ and children affected by LS show developmental delays prior to the onset of regressive phenotypes.^75–77^ Defective mitochondrial function may thus interfere with physiological neurogenesis.^78,79^ Defective embryonic development and delayed neurogenesis have been also observed in LS mice^80,81^ and in brain organoid models of LS.^82,83^ Our previous works suggested that drugs promoting neurogenesis could represent potential therapies for patients with LS.^84,85^ Here, we further confirm the neurodevelopmental impact of mitochondrial defects and demonstrate that the introduction of heteroplasmic mtDNA variants in otherwise healthy stem cells is sufficient to disrupt early neuronal specification.

As heteroplasmy levels decreased upon extensive proliferation of mtDNA edited iPSCs, it is possible that selection mechanisms exist, raising the possibility that treatment strategies might be developed based on harnessing such mechanisms. Changes of heteroplasmy have been already described in iPSC lines upon long term expansion, with some studies reporting increase and dominance^86–88^ and other studies showing a reduced level during culture.^66,67,70^ Shifting heteroplasmy towards more WT copies by specifically targeting mutant mtDNA using mitoTALENs,^89^ mitoZFNs,^90^ or mitoARCUS^91^ has been repeatedly demonstrated *in vitro*^92–95^ and *in vivo*.^96,97^ However, it remains to be determined whether heteroplasmy changes can be actively modulated by hijacking naturally occurring selection mechanisms, such as processes that modify mtDNA copy number, mitochondrial biogenesis, or mitophagy. Future studies performed at single-cell level should tackle these issues to uncover how heteroplasmy level is maintained or selected against in proliferative and postmitotic cells. At the same time, since approximately 20 % heteroplasmy was still eliciting transcriptional reprogramming in iNs, it is possible that treatment strategies simply reducing heteroplasmy would not be sufficiently effective in patients. In fact, unlike CV variants that typically cause LS when present at high levels of heteroplasmy, CI heteroplasmic variants can be pathogenic in patients and iPSC models even at low mutation load.^19,98^

More generally, this work demonstrates that mtDNA base editing could represent a novel tool for disease modeling studies in addition to CRISPR/Cas engineering for nuclear gene defects.^18,25^ DdCBE has been recently employed as a method for correcting pathogenic patient-derived iPSCs carrying the homoplasmic mtDNA variant m.4300A>G in the *MT-TI* gene encoding a tRNA and causative maternally inherited hypertrophic cardiomyopathy.^44^ DdCBE was also applied to human primary adult liver stem cell-derived organoids^99^, as well as a number of different animal models and cell types.^37–43^ To our knowledge, our work is the first report of mtDNA engineering to introduce pathogenic mtDNA variants in iPSCs for precise disease modeling. Future studies are warranted to investigate the presence and significance of nuclear off-targets in more details, as the use of DdCBE was found to be associated with nuclear off-target editing beyond NUMTs.^100,101^ Nonetheless, the ability to precisely modify mtDNA in iPSCs could represent an important advance in human disease modeling. mtDNA edited isogenic iPSC lines may be harnessed in the future to investigate the developmental role of mitochondrial defects or the mechanisms underlying heteroplasmy regulation in human cells.

## ACKNOWLEDGEMENTS

We thank Michal Minczuk (Cambridge, UK) for critical discussion. We acknowledge support from the Deutsche Forschungsgemeinschaft (DFG) (PR1527/13-1, PR1527/14-1, PR1527/15-1, PR1527/18-1 to A.P.; RO5380/1-1, RO5380/10-1, RO5380/12-1, RO5380/14-1 to A.R.), ERDERA and Federal Ministry of Research, Technology and Space (BMFTR) (SynLeigh #01GM2602A to A.P.), the European Commission’s Horizon Europe Programme (SIMPATHIC #101080249 to A.P.), AFM-Téléthon (28545, 29044, 30119, to A.P.; 25179 to A.R.), the patient organizations Mitocon, Cure Mito, and Cure ATP6 (to A.P.), North Rhine-Westphalia (NRW) start-up grant (IN-SU-3-001 to A.R. and J.D.), and the Leibniz Competition (SAW) Cooperative Excellence project (K642/2024 to A.R.).

## AUTHOR CONTRIBUTION STATEMENT

Conceptualization, I.T. A.P.; Methodology, I.T., F.T., D.H., P.L., V.R.B.V.; Formal Analysis, J.D. C.F., A.P.P., A.Z.; Resources, E.M., A.P., A.R., V.C.; Writing –Original Draft, A.P.; Writing – Review & Editing, A.P., I.T., A.R., V.C.; Supervision, A.P., A.R., E.B. V.C.; Visualization, I.T., A.P.; Funding Acquisition, A.P., A.R.

## METHODS

### Ethics

iPSC lines were used in accordance with the ethical approval obtained by the Ethics Committee of the Medical Faculty of Heinrich Heine University (study number 2019-681 approved on October 11, 2019).

### Induced pluripotent stem cells (iPSCs)

The WT inducible NGN2 iPSC line BIHi005-A-24 was generated from the healthy male iPSC line BIHi005-A, which had been reprogrammed from dermal fibroblasts using non-integrating mRNA, by TALEN-mediated insertion of a constitutive m2rtTA cassette and a doxycycline-inducible NGN2 cassette into the two AAVS1 alleles.^102,103^ The cells were routinely monitored for mycoplasma contamination by PCR using nine primers to amplify the six most common mycoplasma strains. The positive control and internal control were kindly provided by Dr. Cord Uphoff (DSMZ, Germany). The cells were cultured on Geltrex (Gibco)-coated plates using StemMACS iPS-Brew XF medium (Miltenyi Biotec), supplemented with MycoZap Plus-CL (Lonza) in a humidified atmosphere of 5 % CO_2_ at 37 °C and 5 % oxygen. They were passaged at 70-80 % confluence with either 0.5 µM EDTA (Invitrogen) in 1 x PBS (Gibco) for routine maintenance or Accutase (Gibco) for single-cell suspensions and replated in complete medium supplemented with the CEPT^104^ cocktail (R&D Systems). Karyotyping was conducted using the Illumina Infinium Global Screening Array (GSAMD-24v3-0-EA_20034606) by Life&Brain.

### HEK cells

HEK293 cells were cultured in Dulbecco’s Modified Eagle Medium GlutaMAX (DMEM GlutaMAX; Gibco) supplemented with 10 % fetal bovine serum (FBS; Gibco), non-essential amino acids (MEM NEAA, Gibco), 1 mM sodium pyruvate (Gibco) and MycoZap Plus-CL (Lonza) at 37 °C in a humidified atmosphere containing 5 % CO_2_. They were passaged using 0.05 % Trypsin-EDTA (Gibco).

### Derivation of induced neurons (iNs)

iNs were obtained as previously described.^102^ Briefly, WT iPSC line BIHi005-A-24 and its mtDNA edited derivatives (ND1.1, ND1.2, ND1.3, ND5.1, ND5.2, ND5.3) were split with Accutase and 3-6 × 10E5 iPSCs per well were seeded in a Geltrex (Gibco, USA) coated 6-well plate (Greiner Bio-One, Austria) using StemMACS iPS-Brew medium supplemented with CEPT cocktail (R&D Systems). On day 0 and day 1, we added induction medium (DMEM-F12, 1x N2 supplement, 1x NEAA, 1 mg/ml Pen/Strep [all Gibco, USA], 10 ng/ml human NT-3 [PeproTech, USA], 10 ng/ml human BDNF [Miltenyi Biotec, Germany], 0.2 μg/ml murine laminin [Sigma-Aldrich, USA]) freshly supplemented with 3 μg/ml doxycycline (dox) (Sigma-Aldrich, USA). On day 2, medium was changed to neuronal medium (Neurobasal, 1x B-27 supplement, 1x GlutaMAX, 1 mg/ml Pen/Strep [all Gibco, USA], 1x MycoZapTM Plus-CL (Lonza, Switzerland), 10 ng/ml human NT-3 [PeproTech, USA], 10 ng/ml human BDNF [Miltenyi Biotec, Germany], 0.2 μg/ml murine laminin [Sigma-Aldrich, USA]) freshly supplemented with 3 μg/ml dox (Sigma-Aldrich, USA). Medium was exchanged daily until day 4. From day 6 on, the medium was replaced every other day with neuronal medium freshly supplemented with 3 μg/ml dox (Sigma-Aldrich, USA) and 2 μM AraC (Sigma-Aldrich, USA) until day 9 when media containing AraC were removed. From that day forward half media changes were performed every second day until the cells were harvested for subsequent experiments.

### mtDNA base editing

For generation of the DdCBE plasmids required for mtDNA editing, TALE sequences were designed using TALEN Targeter (https://talent.cac.cornell.edu/node/add/talen) and Paired Target Finder (https://talent.cac.cornell.edu/node/add/talef-off-paired). The TALE sequences were ordered as gene synthesis from GeneArt (Thermo Fisher Scientific) and codon optimized for human hosts. They were cloned into Addgene plasmids #158096 and #158092 via golden gate cloning using Esp3I (NEB) and fluorescence marker (mCherry, mNeonGreen) were added to the end of the constructs linked via T2A sequences. The plasmids were verified via Sanger sequencing.

WT iPSCs BIHi005-A-24 were transfected either with only the H-strand DdCBE as mock-treated control, or both DdCBE plasmids using ScreenFect A-Plus according to manufacturer’s instructions. Briefly, 1.2 µl ScreenFect A-Plus reagent were mixed with 300 ng of each plasmid (600 ng total pDNA) and incubated for at least 20 min at 22 °C. The ScreenFect/plasmid emulsion was mixed with 8 x 10E5 singularized iPSCs and plated in one well of a 24-well plate in media without MycoZap. After 24 h, media were replaced with standard cultivation media containing MycoZap. At 48 h post-transfection, cells positive for both mNeonGreen and mCherry were sorted by FACS (BD FACS AriaIII, Core Facility Flow Cytometry) and seeded in low density (300 and 500 cells/well of a 6-well plate) to allow for growth of single colonies. Approximately 11-13 days after seeding, single colonies were picked and partly used for further cultivation, the other part used for an initial screen via Sanger sequencing (Eurofins). For Sanger sequencing analysis, the cells were lysed using Phire Animal Tissue Direct PCR Kit (Thermo Fisher Scientific) following the dilution protocol. Lysates were used as template in a PCR reaction to amplify the target region with Q5 High-Fidelity DNA Polymerase (NEB) resulting in amplicons of about 1 kb size. For estimation of editing efficiency, Sanger traces were analyzed using EditR (https://moriaritylab.shinyapps.io/editr_v10/).^105^

### Next generation sequencing (NGS) of whole mtDNA

Whole mtDNA sequencing was performed after PCR amplification with two sets of primers (MT1F: CCCATAACACTTGGGGGTAG/ MT1R: TAGAAGTGTGAAAACGTAGGCTTG and MT2F: GCCCTAGCCCACTTCTTACC/ MT2R: TGTAAGTTGGGTGCTTTGTGTT) using the PrimeSTAR Max DNA Polymerase (Takara Bio). A library was constructed from purified amplicons using xGen DNA Library Prep Kit (IDT) and then sequenced on a MiSeq instrument (Illumina) with 150 bp paired-end reads. The sequences were aligned with SAMtools on the rCRS (NC_012920.1) reference^106^ and filtered through the MitoScape software.^107^ mtDNA variants were called combining Mutect2 and mtDNA-Server 2 tools’ outputs.^108–110^ All G>A and C>T substitutions arising from deaminase treatment were systematically evaluated. No heteroplasmy threshold was applied during variant filtering, with the aim of capturing low-frequency variants potentially undergoing clonal expansion. Identified variants were manually curated and classified according to their predicted pathogenic potential. Emphasis was placed on missense variants classified at least as VUS− according to the APOGEE2 predictor,^111^ prioritizing those with a plausible deleterious functional impact. For variants located within mitochondrial tRNA genes, pathogenicity was inferred by integrating MitoTIP scores with population frequency data derived from healthy cohorts.^112^ No classification was performed for variants located within non-coding regions.

### mtDNA sequencing with Mitopore

Mitochondrial DNA (mtDNA) sequencing and analysis were performed using the previously established Mitopore workflow.^46^ Briefly, the mitochondrial genome was amplified by PCR using two overlapping amplicons of approximately 9 kb, collectively covering the entire mtDNA. PCR amplification was performed for 30 cycles using Q5 High-Fidelity DNA Polymerase (NEB) to minimize amplification bias. Amplicons were purified, pooled per sample, and subjected to library preparation using the Oxford Nanopore Technologies (ONT) ligation sequencing and native barcoding workflow. Barcoded samples were pooled and sequenced on a Flongle flow cell using a MinION Mk1C sequencing device (Oxford Nanopore Technologies). Resulting FASTQ files were analyzed using Mitopore as previously described.^46^ Briefly, sequencing reads were aligned to the revised Cambridge Reference Sequence (rCRS; NC_012920) using Minimap2, followed by mitochondrial variant calling using Mutserve. Variant annotation and visualization were performed within the Mitopore analysis workflow.

### Assessment of mitochondrial membrane potential (MMP)

For MMP quantification, we applied a high-content analysis (HCA)-based live-cell detection assay that we previously established.^113^ Briefly, we seeded iPSCs or iNs onto black-wall, clear-bottom 96-well plates (SCREENSTAR, Greiner) pre-coated with Geltrex (Thermo Fisher Scientific) at a density of 2 x 10E4 cells/cm^2^ and incubated them in their respective media overnight at 37 °C and 5 % CO_2_. On the day of the assay, we live-stained the cells with 10 nM tetramethylrhodamine methyl ester (TMRM) (Molecular Probes, Life Technologies) and 1 µg/ml Hoechst 33342 (Thermo Fisher Scientific) in the appropriate media for 30 min at 37 °C and 5 % CO_2_. After 30 min incubation, cells were washed with 100 µl media without phenol red. Cells were kept in 100 µl medium (w/o phenol red) for the duration of the assay. For confirming MMP depolarizing effect, we used the mitochondrial uncoupler carbonyl cyanide-p-trifluoromethoxyphenylhydrazone (FCCP) in combination with the mitochondrial complex III inhibitor antimycin A (AA). Imaging was performed with Operetta CLS high content imaging system (Revvity) with 20x water objective in confocal mode. Imaging time did not exceed 30 min. TMRM and Hoechst 33342 intensity within the cells was analyzed with CellProfiler (version 4.2.5). MMP values are presented as the TMRM intensity in each sample over the TMRM intensity in the WT sample.

### Mitochondrial complex I (CI) enzymatic activity

Biochemical analyses were performed on iPSC cell lysates following the protocol described before.^114^ Briefly, 1 x 10E6 iPSCs were suspended in 0.4 ml of 20 mM potassium phosphate buffer pH 7.5. The iPSC suspensions were homogenized by using 50 µl Hamilton syringe until a homogeneous solution was obtained. iPSC cell lysates were snap-frozen in dry ice and thawed at 37 °C three times. Protein content was measured using BCA assay (ThermoFisher, #23227) and the mitochondrial complex I (CI) and citrate synthase (CS) enzymatic activities were measured spectrophotometrically in 96 well-plates (Molecular Devices, SpectraMax ID5e) as previously described^115^ with some modifications. CI activity (NADH:ubiquinone oxidoreductase) was measured through a direct method by following the oxidation of NADH. iPSC lysates were resuspended in the reaction mix containing 20 mM potassium phosphate buffer pH 8, 0.2 mM NADH (Sigma #N4505), 1 mM sodium azide, 0.1 % BSA, 1 mM EDTA Na_4_ pH 7.4. After pre-incubation at 30 °C, the CI reaction was started with 50 μM CoQ_1_ (Cayman Chemical, #18741) and then inhibited with 5 μM rotenone (Sigma, #R8875). Both CoQ_1_ and rotenone reactions were recorded for 2 min at λ=340 nm, 30 °C. Specific CI activity corresponded to the subtraction of Δabsorbance CoQ1/min and Δabsorbance Rotenone/min. CS activity was measured using an indirect method by following the formation of thionitrobenzoate (TNB) derivative of DTNB (Sigma, #D8130). iPSC lysates were resuspended in the mix containing 100 mM This-HCl pH 8, 0.2 % Triton X100, 100 µM DTNB, 300 µM Acetyl-CoA (Sigma, #A2181) and incubated for 2 min at 30 °C. 500 µM oxaloacetate (Sigma, #O4126) was added to start the CS reaction and recorded at λ= 412 nm for 2 min at 30 °C. CS activity was used to normalize the CI activity assays. The enzymatic specific activity for CI and CS was expressed as amount of substrate converted per time unit per mass unit of cell lysates (nmol. min^−1^ x mg^−1^).

### Targeted metabolomics profiling

iPSCs were washed with ice cold PBS and detached using accutase at 37 °C for 7 min. The reaction was stopped with cold PBS and cells kept on ice during quantification using an automatic cell counter. 1 x 10E6 iPSCs/sample were pelleted using a tabletop centrifuge at 500 x g for 5 min at 4 °C. The supernatant was discarded and the pellet was snap-frozen in liquid nitrogen and stored at −80 °C until measurement. Extraction of the targeted metabolites was performed by adding 300 µl Methanol/H_2_O (80/20; v/v) to the 1 x 10E6 cells, which were homogenized by ceramic beads CK14 using Precellys lysing kit. The corresponding isotopically labelled internal standards were added to the extraction solution. The targeted compounds were analyzed by UPLC-MS/MS. The system consists of a UPLC I-Class (Waters) coupled to a tandem mass spectrometer Xevo-TQ-XS (Waters). Electrospray ionization was performed in the negative ionization mode for the compounds fructose-1,6-bishphosphate, phosphoenolpyruvate (PEP) and 2/3-phosphoglycerate.^116^ The TCA, energy metabolites (AMP, ADP and ATP) as well acetyl-and succinyl-CoA were measured in the positive ionization mode.^117–119^ Mass spectrometric quantitation of the compounds was carried out in the multiple reaction monitoring (MRM) mode. MassLynx software (v. 4.2) was used for instrument control and data acquisition. Quantitation analysis was performed by TargetLynx XS software.

### RNA sequencing of mtDNA edited iNs

Bulk RNA-sequencing of iNs was carried out for the following samples: WT, ND1.1, ND1.2, ND1.3, ND5.1, ND5.2, ND5.3. Each experimental group was represented by three biological replicates obtained from independent differentiation experiments (n = 21 samples in total). Total RNA was isolated using the RNeasy kit (QIAGEN), and RNA quality and integrity were assessed using an Agilent TapeStation system (Agilent Technologies). Briefly, total RNA was mixed with 1 μg of a DNA oligonucleotide pool comprising 50-nt oligonucleotides complementary to the entire length of each rRNA (28S, 18S, 16S, 5.8S, 5S, and 12S rRNA), incubated with 1 U of RNase H (Hybridase Thermostable RNase H, Epicentre), purified using RNA Cleanup XP beads (Agencourt), treated with TURBO DNase according to the manufacturer’s rigorous treatment protocol (Thermo Fisher Scientific), and purified again using RNA Cleanup XP beads. The resulting rRNA-depleted RNA was fragmented and processed into strand-specific cDNA libraries using the TruSeq Stranded Total RNA LT Sample Prep Kit (Illumina). Libraries were sequenced on an Illumina NovaSeq 6000 platform using paired-end sequencing (2 × 150 bp), generating approximately 40–50 million reads per sample.

### RNA sequencing data processing and quantification

Raw paired-end RNA-sequencing reads were assessed using FastQC (v0.12.0) and aligned to the human reference genome (GRCh38, Ensembl release 109) using STAR (v2.7.10a) in two-pass mode. Gene-level read counts were generated during alignment using the STAR --quantMode GeneCounts option. Raw count matrices were used for downstream differential expression analysis. Differential expression analysis was performed in R (v4.3.2) using DESeq2 (v1.42.0). Genes with low expression were excluded prior to analysis by retaining genes with >1 count per million (CPM) in at least three biological replicates. DESeq2 normalization and dispersion estimation were performed using default settings. Differential expression was assessed using negative binomial generalized linear models and Wald tests. Pairwise comparisons were performed between individual *MT-ND5* mutant clones (clones ND5.1, ND5.2 and ND5.3) or *MT-ND1* mutant clones (clones ND1.1 and ND1.3) and the parental BIHi005-A-24 control. P values were corrected for multiple testing using the Benjamini–Hochberg procedure. Genes with an adjusted P value <0.05 and an absolute log2 fold change ≥1 were considered differentially expressed. Data processing and visualization were performed in R (v4.3.2). Heatmaps were generated using pheatmap (v1.0.12). Normalized expression values were log2-transformed and standardized by gene using Z-score transformation. Hierarchical clustering of genes was performed using Euclidean distance and complete linkage, while sample order was fixed where indicated to facilitate comparison between mutant clones and controls. Data analysis and visualization were performed using packages from the tidyverse suite (v2.0.0), including dplyr (v1.1.4) and ggplot2 (v3.4.4).

### Mitochondrial heteroplasmy analysis

Heteroplasmy at the targeted *MT-ND1* and *MT-ND5* loci was quantified from reads mapping to the mitochondrial genome (revised Cambridge Reference Sequence, NC_012920.1). PCR and optical duplicates were identified and removed using Picard Tools (v2.27.5). To minimize potential interference from NUMTs, only uniquely mapped reads with a mapping quality score ≥30 and base quality ≥30 at the variant position were considered. Nucleotide counts and sequencing depth at the targeted positions were determined using samtools mpileup (v1.17). Heteroplasmy was calculated as the proportion of high-quality reads carrying the variant allele relative to the total number of reads covering the corresponding position and expressed as a percentage. Clones carrying the targeted mtDNA variants were retained for phenotypic and transcriptomic analyses. For clone ND1.2 the intended mutation was not detected and was therefore excluded from mutant-versus-control comparisons.

### Statistical analysis

Data are expressed as mean and standard deviation (mean ± SD) where normality of the distribution could be verified, or as median and quartiles (median [25^th^;75^th^ percentiles]) otherwise. Significance was assessed using parametric tests (Brown-Forsythe and Welch ANOVA with Dunnet’s T3 multiple comparison) for normally distributed data and non-parametric tests (Kruskal-Wallis with Dunn’s multiple comparison) when normal distribution could not be verified. Unless otherwise indicated, data were analyzed and shown using GraphPad-Prism software (v. 10.6.1) (GraphPad Software, USA) or R (v. 4.3.2). Figures were prepared with Adobe Illustrator (v. 30.8.1). Schematic cartoons were made using Biorender.

