## Supplemental figures for "Mitochondrial genome editing reveals the impact of complex I heteroplasmic variants on human neurogenesis"

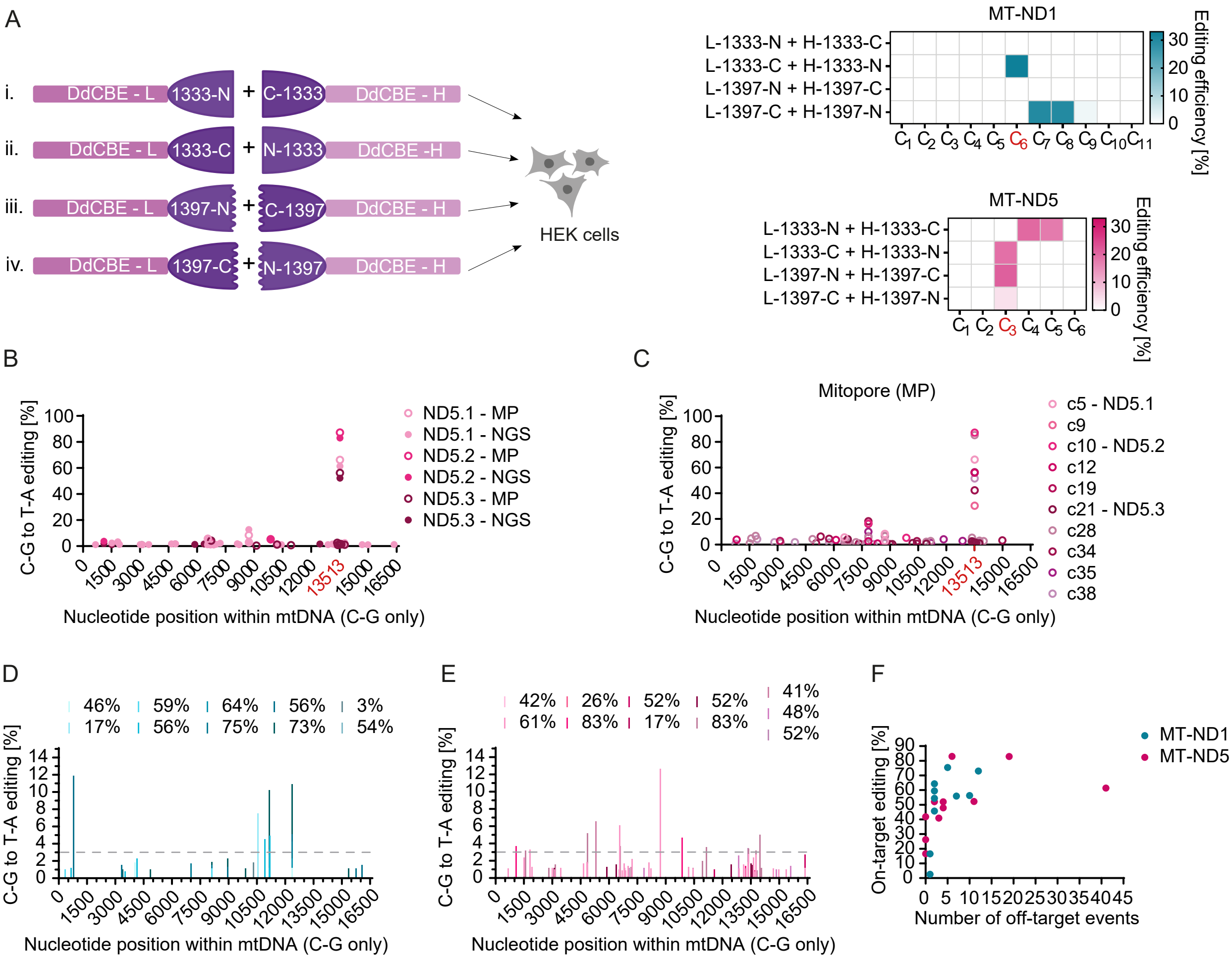

**Figure S1 I Optimization of DddA-derived cytosine base editor (DdCBE) pairs and whole mtDNA sequencing.**

**(A)** Selection of DdCBE split variant and orientation for each target site was identified based on editing efficiency in HEK293 cells. Left: Schematic overview of tested DdCBE orientations. Right: Editing efficiencies of different DdCBE orientations at *MT-ND1* and *MT-ND5* target sites. Targeted cytosines are highlighted in red. Color shades reflect the mean of n=3 independent biological replicates. **(B)** Comparison of whole mtDNA sequencing using Illumina next generation sequencing (NGS) and Mitopore (MP) for the three selected *MT-ND5* edited lines. **(C)** Whole mtDNA sequencing of all *MT-ND5* edited iPSC lines analyzed using Mitopore. **(D-E)** mtDNA-wide off-target events of all *MT-ND1* (D) and *MT-ND5* (E) edited lines analyzed via NGS. Corresponding on-target editing efficiency is indicated in the legend and a dashed line is drawn at 3% for better clarity. **(F)** On-target editing efficiency plotted against number of mtDNA-wide off target events. Results are from NGS analysis.

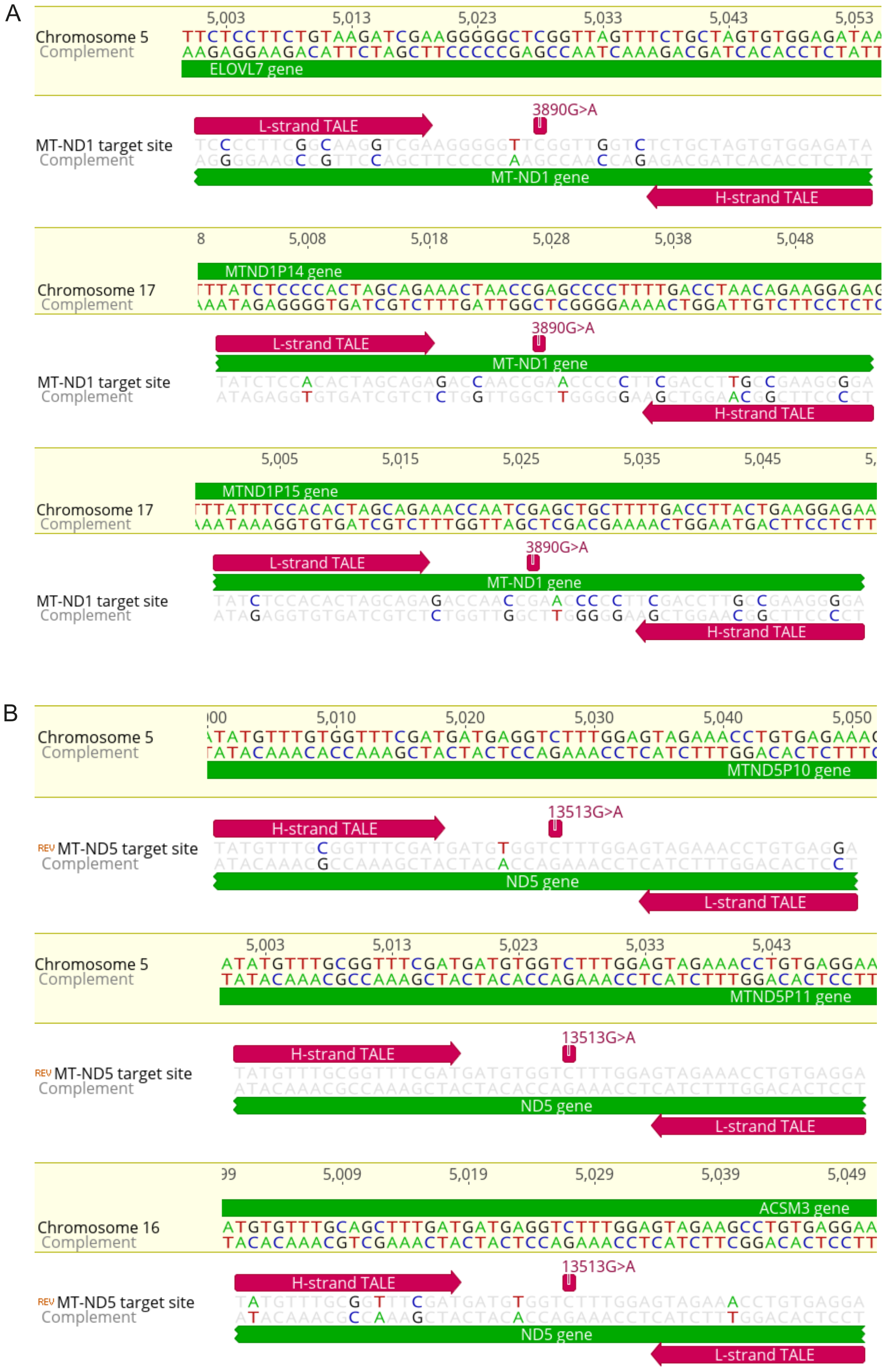


**Figure S2 I Targeted analysis of nuclear off-target editing in mtDNA edited iPSCs.**

**(A-B)** The nuclear genome of edited iPSC lines was screened for putative binding sites of the DdCBE constructs used in this study. Three nuclear regions with the highest predicted binding affinity were selected for each TALE pair [(A) *MT-ND1*, (B) *MT-ND5*]. Bases where the mitochondrial target site differ from its nuclear counterpart are highlighted in color. No nuclear off-target editing could be detected at either of the 6 sequenced pseudogenes.

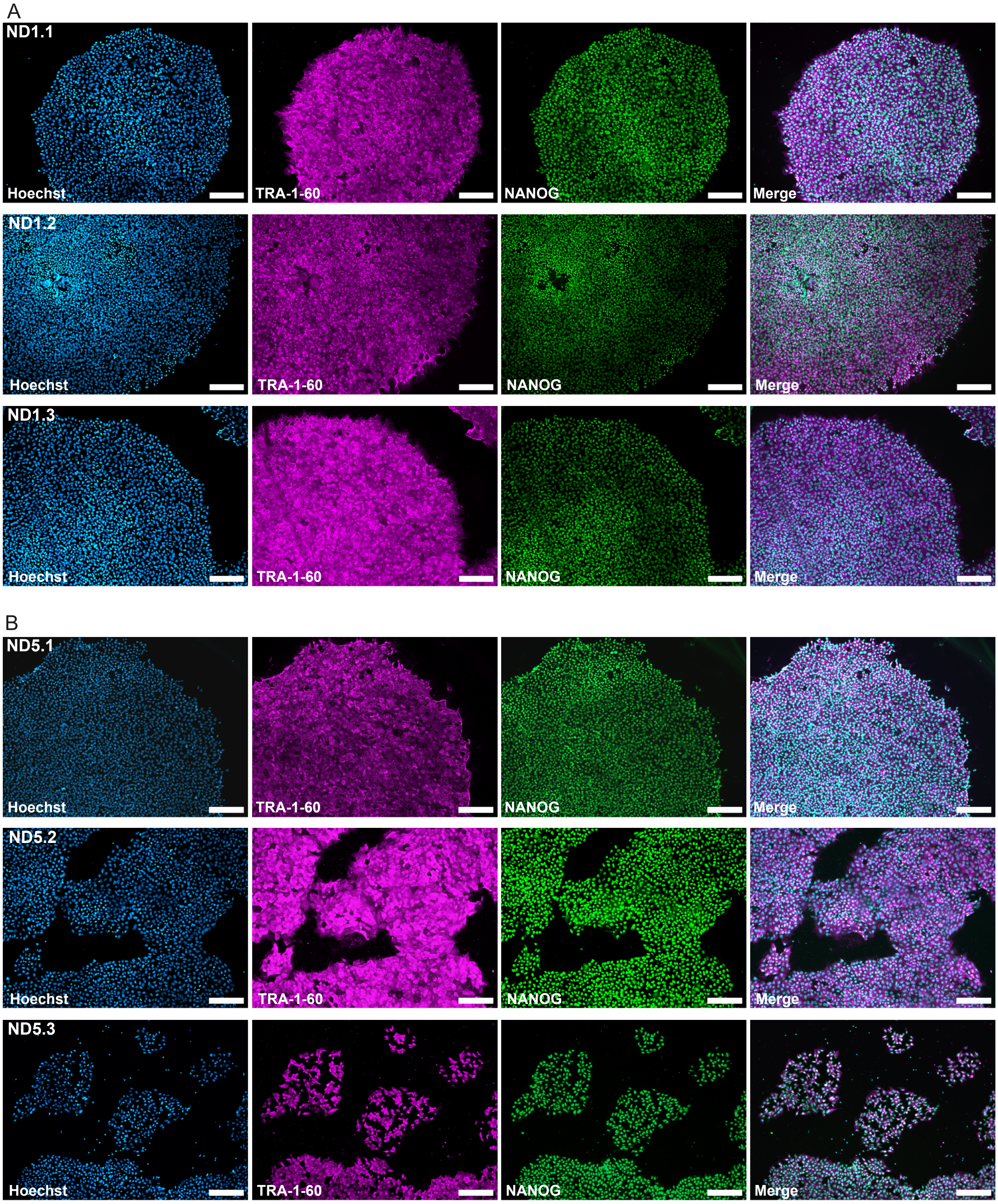


**Figure S3 I mtDNA edited iPSC lines retain pluripotency.**

**(A-B)** All selected edited lines for *MT-ND1* (A) and *MT-ND5* (B) retained the expression of pluripotency-associated protein markers TRA-1-60 (magenta) and NANOG (green). Scale bars represent 200 µm.

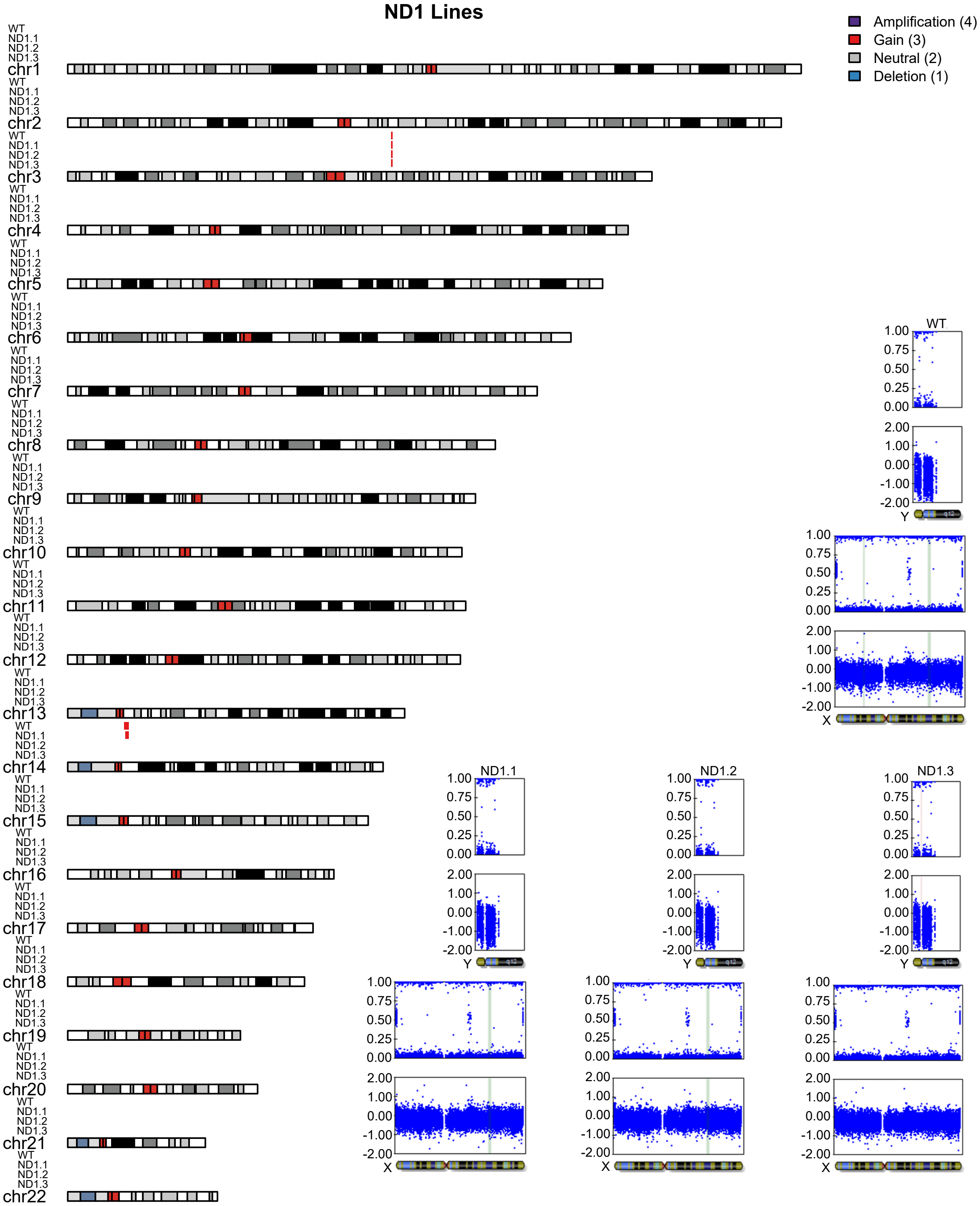


**Figure S4 I *MT-ND1* edited iPSC lines retain normal karyotype.**

The karyotypes of *MT-ND1* edited iPSC lines (ND1.1, ND1.2, ND1.3) were compared to that of the isogenic wildtype iPSC line (WT). A small gain on chromosome 3 was present in all lines and interpreted as a shared line-specific copy number variation (CNV). Apparent alterations involving chromosome 14 were considered likely array-related artifacts (including short-arm signal abnormalities). Calls on chromosome X were attributed to sex chromosome normalization effects in this male line and were not considered true copy number losses. No clone-specific pathogenic CNVs were identified.

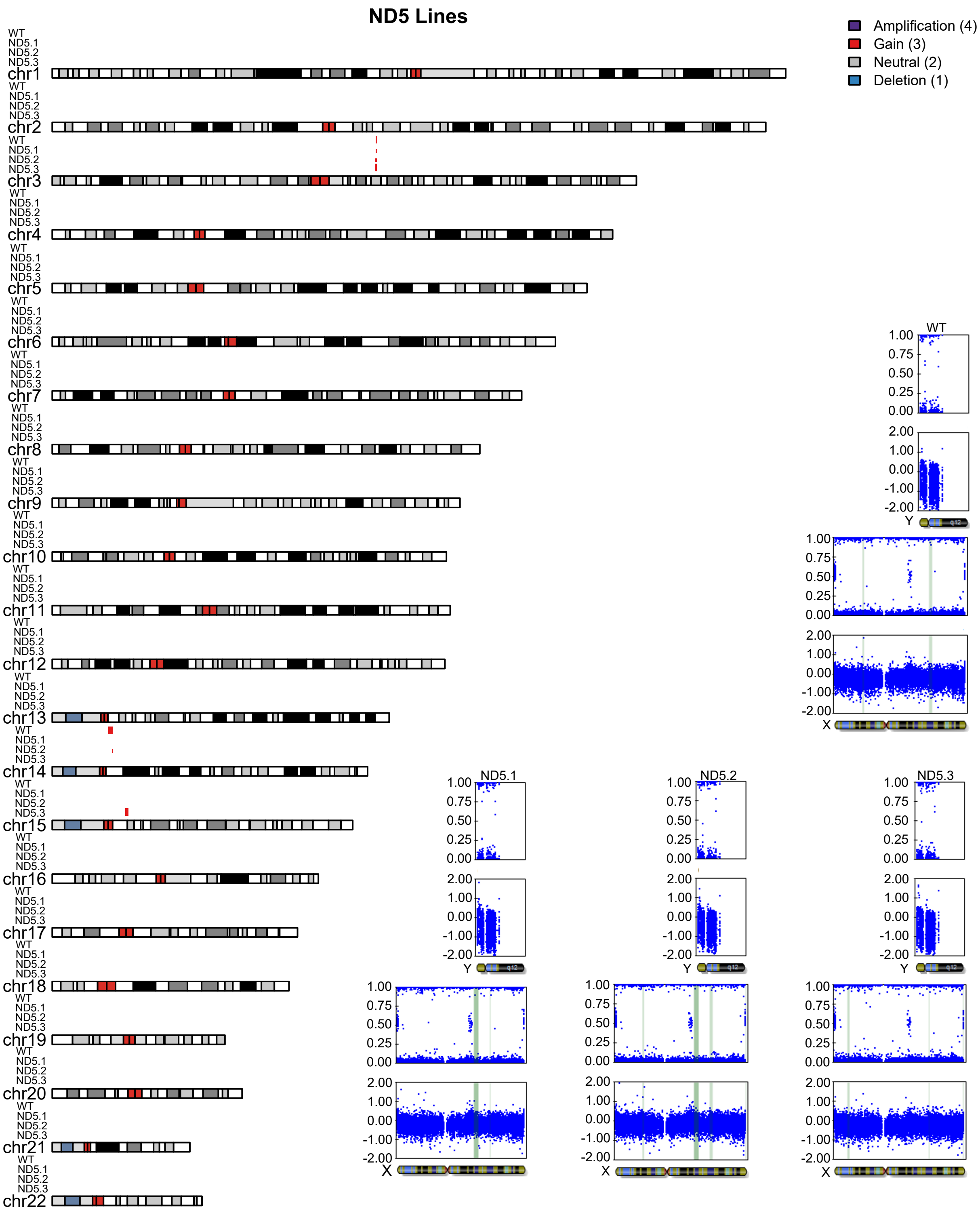


**Figure S5 I *MT-ND5* edited iPSC lines retain normal karyotype.**

The karyotypes of *MT-ND5* edited iPSC lines (ND5.1, ND5.2, ND5.3) were compared to that of the isogenic wildtype iPSC line (WT). A small gain on chromosome 3 was present in all lines and interpreted as a shared line-specific CNV. Apparent alterations involving chromosomes 14 and 15 were considered likely array-related artifacts (including short-arm signal abnormalities). Calls on chromosome X were attributed to sex chromosome normalization effects in this male line and were not considered true copy number losses. No clone-specific pathogenic CNVs were identified.

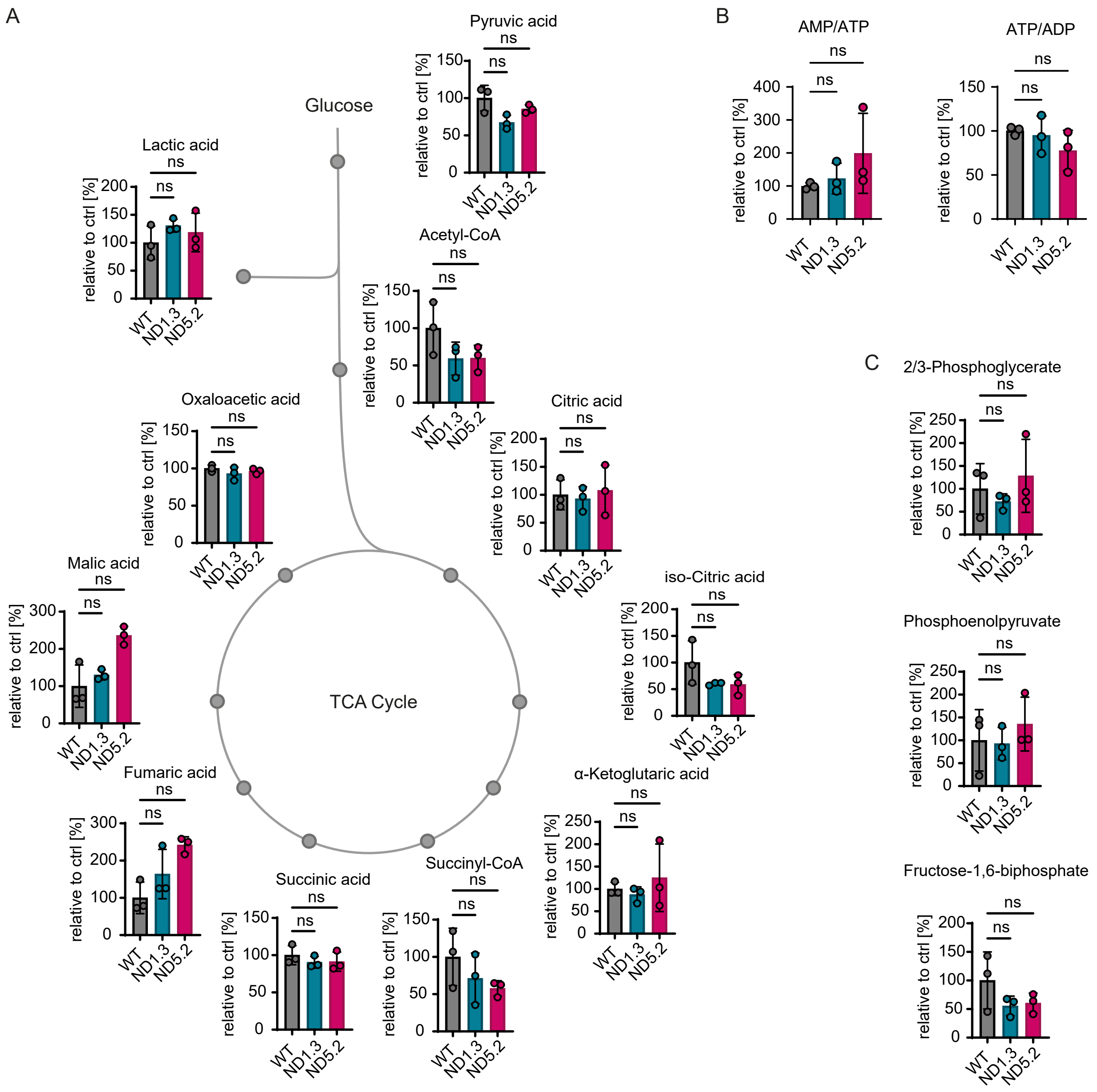


**Figure S6 I Targeted metabolomic profiling in mtDNA edited iPSCs.**

Targeted metabolomics for TCA cycle metabolites **(A)**, ATP amount **(B)**, and glycolysis-related metabolites **(C)** in mtDNA edited iPSC lines compared to isogenic wildtype iPSCs (WT). Values and error bars reflect the mean ± s.d. of n=3 independent biological replicates; ns: not significant; Brown-Forsythe and Welch ANOVA with Dunnett's T3 multiple comparison or Kruskal-Wallis test with Dunn's multiple comparison where normality test failed.

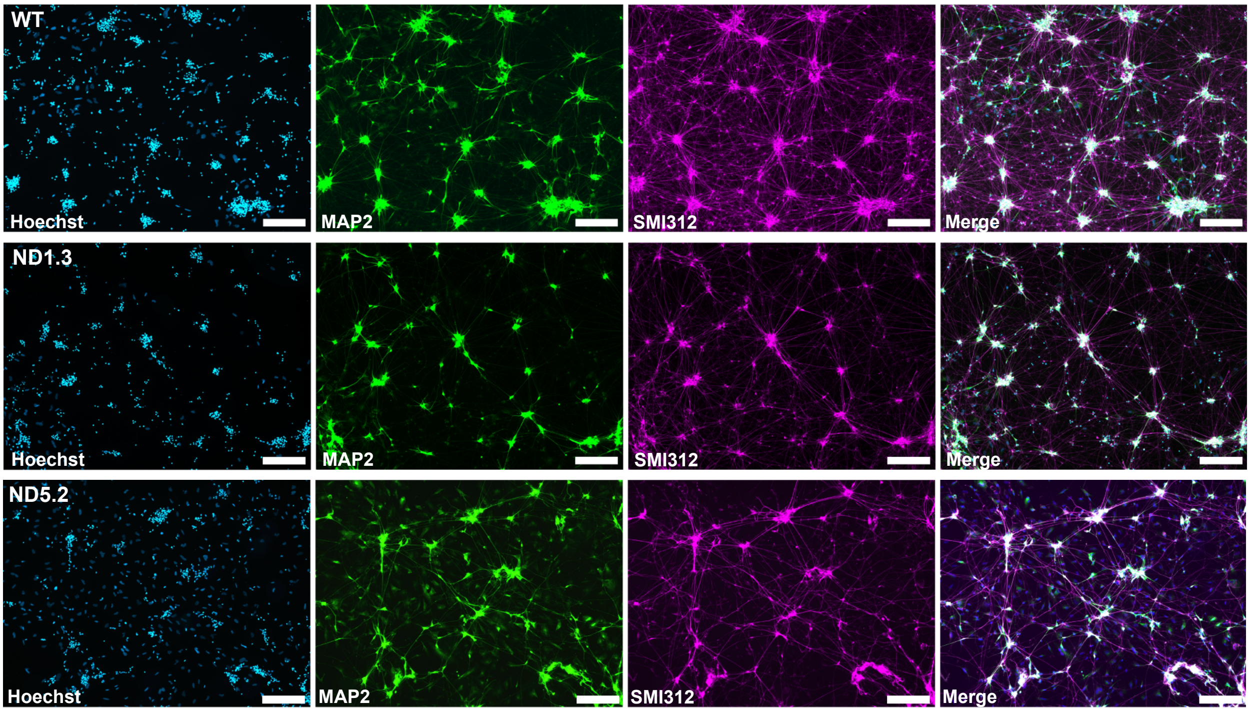


**Figure S7 I Generated iNs express neuronal markers.**

After 14 days of differentiation, WT, ND1.3, and ND5.2 iNs were stained against the neuronal markers MAP2 and SMI312, confirming neural identity as well as development of dendritic and axonal structures, respectively. Scale bars represent 200 µm.

### Supplemental Table Legends

**Table S1 I NGS whole mtDNA analysis of *MT-ND1* edited cells.**

Sheet 1: all samples. Sheet 2: Calculated average off-target editing of mock treated control, bulk sample, ND1.1, ND1.2, and ND1.3. Sheet 3: overview of all 10 edited lines.

**Table S2 I NGS whole mtDNA analysis of *MT-ND5* edited cells.**

Sheet 1: all samples. Sheet 2: Calculated average off-target editing of mock treated control, bulk sample, ND5.1, ND5.2, and ND5.3. Sheet 3: overview of all 11 edited lines. Sheets 4 and 5: ND5.2 and ND5.3 with their subclones.

**Table S3 I Mitopore whole mtDNA analysis of *MT-ND5* edited cells.**

Sheet 1: all samples. Sheet 2: comparison to Illumina NGS.

**Table S4 I Differential gene expression (DGE) analysis with SD, fold change and GO pathways of iNs after 14 days of differentiation.**
